# Synergistic protection against secondary pneumococcal infection by human monoclonal antibodies targeting distinct epitopes

**DOI:** 10.1101/2022.04.20.488872

**Authors:** Aaron D. Gingerich, Fredejah Royer, Anna L. McCormick, Anna Scasny, Jorge E. Vidal, Jarrod J. Mousa

**Affiliations:** Center for Vaccines and Immunology, College of Veterinary Medicine, University of Georgia, Athens, GA, USA; Department of Infectious Diseases, College of Veterinary Medicine, University of Georgia, Athens, GA, USA; Department of Microbiology and Immunology, University of Mississippi Medical Center, Jackson, Mississippi, USA; Department of Biochemistry and Molecular Biology, Franklin College of Arts and Sciences, University of Georgia, Athens, GA, USA

## Abstract

*Streptococcus pneumoniae* persists as a leading cause of bacterial pneumonia despite the widespread use of polysaccharide-based vaccines. The limited serotype coverage of current vaccines has led to increased incidence of non-vaccine serotypes, as well as an increase in antibiotic resistance among these serotypes. Pneumococcal infection often follows a primary viral infection such as influenza virus, which hinders host defense and results in bacterial spread to the lungs. We previously isolated human monoclonal antibodies (mAbs) against the conserved surface antigen pneumococcal histidine triad protein D (PhtD), and we demonstrated that mAbs to this antigen are protective against lethal pneumococcal challenge prophylactically and therapeutically. In this study, we elucidated the mechanism of protection of a protective anti-pneumococcal human mAb, PhtD3, which is mediated by the presence of complement and macrophages in a mouse model of pneumococcal infection. Treatment with mAb PhtD3 reduced blood and lung bacterial burden in mice, and mAb PhtD3 is able to bind to bacteria in the presence of the capsular polysaccharide, indicating exposure of surface PhtD on encapsulated bacteria. In a mouse model of secondary pneumococcal infection, protection mediated by mAb PhtD3 and another mAb targeting a different epitope, PhtD7, was reduced, however, robust protection was restored by combining mAb PhtD3 with mAb PhtD7, indicating a synergistic effect. Overall, these studies provide new insights into anti-pneumococcal mAb protection and demonstrate for the first time that mAbs to pneumococcal surface proteins can protect against secondary pneumococcal infection in the mouse model.

**Author Summary:** The persistence of Streptococcus pneumoniae as a leading cause of bacterial pneumonia despite numerous approved pneumococcal vaccines is a serious threat to public health globally. Currently, prophylactic and therapeutic options for *Streptococcus pneumoniae* are constrained by the limited serotype coverage of vaccines and the emergence of antibiotic resistant strains. An additional hurdle to overcome is the incidence of secondary pneumococcal infection following a viral infection, which leads to increased mortality. Here, we determined the mechanism of action of a monoclonal antibody (mAb) that targets *Streptococcus pneumoniae*. We found that mAb PhtD3 operates through macrophage and complement mediated functions. mAb PhtD3 was also discovered to reduce bacterial titers in the lungs and blood and bind to a related antigen PhtE. We also tested additional mAbs and discovered that two unique mAbs to the antigen PhtD conferred protection in a pneumococcal-influenza virus co-infection model. Our study provides new insights into the mechanisms and therapeutic potential of mAbs targeting conserved proteins of *Streptococcus pneumoniae*.

## Introduction

*Streptococcus pneumoniae* is designated a priority pathogen by the World Health Organization due to widespread infections and high mortality nearing 1 million deaths worldwide each year (1). Currently, pneumococcal vaccination for disease prevention and antibiotics for disease treatment are widely used, and have been highly successful in reducing the morbidity and mortality associated with pneumococcal infection. However, several factors remain to be addressed that have resulted in persistent rates of infection worldwide, including access and acceptance of pneumococcal vaccines across the globe (2). Pneumococcal vaccines elicit antibodies targeting the capsular polysaccharide (CPS), which is an important virulence factor of most pneumococcal strains. However, at least 100 distinct CPS serotypes exist (3), and current vaccines only target a fraction of these, with a 23-valent pneumococcal polysaccharide vaccine (PPSV23), or multivalent pneumococcal conjugate vaccines (PCV7, PCV10, PCV13, PCV15 or PCV20) available. Due to the limited coverage of these vaccines, there has been an emergence of non-vaccine serotypes and nonencapsulated strains that are unaffected by the vaccine (4). An additional hurdle to overcome is antibiotic resistance, which is common among non-vaccine serotypes, with some isolates being multidrug resistant (5). Vaccine efficacy is also variable among each included pneumococcal serotype. For example, pneumococcal serotype 3 is a major cause of invasive pneumococcal disease in children and adults despite its inclusion in the PCV13 vaccine (6–10). This is likely due to the serotype 3 CPS being poorly immunogenic with a thick CPS (11), and the copious amount of CPS shedding (12).

Pneumococcal disease has higher incidence following influenza infection, significantly increasing the risk of severe disease resulting in hospitalization and death (13). During the 2009 H1N1 influenza pandemic, an estimated 29-55% of fatalities occurred in patients with bacterial co- infections (14, 15). Additionally, secondary pneumococcal infection is hypothesized to have caused most deaths during the 1918 influenza pandemic (16). Influenza infection initiates inflammation in the airways leading to damage of epithelial cells, which provides an increase in nutrients (17) and attachment sites for *S. pneumoniae* (18, 19). Influenza virus alters mucus production and decreases ciliary action, reducing mucociliary clearance of *S. pneumoniae* (20). The innate immune response following influenza infection is also impaired. Alveolar macrophage-mediated clearance plays a key role in bacterial defense (18), yet influenza infection depletes alveolar macrophages, thus impairing early bacterial clearance (21, 22). Neutrophil impairment has also been shown to occur during influenza infections (23). Furthermore, in the mouse model antibiotic treatments are ineffective at preventing mortality in co-infections (24).

Monoclonal antibodies (mAbs) are a promising tool for broad treatment against numerous pneumococcal serotypes while simultaneously avoiding the complications of drug resistance (25). We previously isolated the first human monoclonal antibodies to pneumococcal protein antigens, including the pneumococcal histidine triad protein (PhtD) and pneumococcal surface protein A (PspA) (26). mAbs to PhtD prolonged survival of pneumococcal infected mice when administered before or after infection (26). While *in vitro* opsonophagocytic assays showed anti-PhtD mAbs lead to an increase in phagocytic activity, the mechanism of protection *in vivo* remains undefined. In this study we defined the mechanistic correlation of protection for a human anti-PhtD mAb, PhtD3, and determined that protection is mediated by macrophages and complement in an intranasal infection model of pneumococcal infection. We also found that bacterial titers in the lungs and blood of infected mice were significantly reduced in mAb PhtD3 treated mouse groups. Additionally, we discovered that a subset of mAbs to PhtD cross-react with the related pneumococcal histidine triad protein E, and this cross-reactivity was correlated to epitope specificity. Finally, we examined the efficacy of anti-PhtD mAbs in a model of secondary pneumococcal infection and found that individual protective mAbs have reduced efficacy in this model, however, administration of two mAbs targeting nonoverlapping epitopes restores protective efficacy.

## Materials and Methods

### Ethics statement

This study was approved by the University of Georgia Institutional Review Board as STUDY00005127. Healthy human donors were recruited to the University of Georgia Clinical and Translational Research Unit, and written informed consent was obtained. Human subjects were recruited for a single blood draw. All animal studies performed were in accordance with protocols approved by the Institutional Animal Care and Use Committee of the University of Georgia.

### Bacterial strains and growth conditions

Bacterial colonies were grown on BD Trypticase Soy Agar II with 5% Sheep Blood (BD, Franklin Lakes NJ). Pneumococcal strains were grown at 37 °C in 5% CO2 in Todd-Hewitt broth (BD, Franklin Lakes NJ) supplemented with 0.5% yeast extract for 12 hrs. Cultures were frozen and stored -80 °C with 10% glycerol until used, and following thawing cultures were washed twice with PBS before being used in experiments. The numbers of CFUs per milliliter of these stocks were determined by plating a single quick-thawed diluted aliquot on sheep’s blood agar plates. The calculated number of CFUs was subsequently used to make dilutions for experiments from aliquots thawed at later times. In each experiment, the actual number of CFUs administered was determined by plating on blood agar at the time of the assay.

### Antibody generation with CHO cells

mAbs PhtD3 and PhtD7 were generated as previously described in Expi293F cells or from hybridoma cultures (26). Alternatively, mAb PhD3 was generated by transient transfection of heavy and light chain plasmids into ExpiCHO cells. ExpiCHO cells were grown in FreeStyle CHO media supplemented with L-glutamine. For transfections, cells were transferred to ExpiCHO media and transfected using the high-titer protocol according to the manufacturer’s instructions.

mAbs were purified from culture and hybridoma supernatants using protein G columns as previously described (26).

### Primary pneumococcal challenge studies

For primary pneumococcal infection studies, 5-7 week old C57BL/6 mice (Charles River) were intranasally infected with pneumococcal serotype 3 strain WU2 by anesthetizing mice via inhalation of 5% isoflurane and administering 40 µL of 5x10^6^ colony-forming units (CFUs) of bacteria in phosphate buffered saline (Corning). Either 2 hrs prior to infection (for prophylaxis studies) or 24 hrs after infection (for therapeutic studies) mice were intraperitoneally inoculated with 15 mg/kg of mAb. Mice were weighed and assessed daily, and mice were euthanized when >20% (for prophylaxis studies) or >30% (for therapeutic studies) of pre-infection body weight was lost, were nonresponsive to manual stimulation, and/or were exhibiting respiratory distress.

### Depletion Studies

Neutrophils were depleted in mice as previously described (27). Briefly, 5-7 week old C57BL/6 mice (Charles River) were intraperitoneally injected with 25 µg of RB6-8C5 antibody 1 day prior to pneumococcal infection. Mice were intraperitoneally inoculated with 15 mg/kg of mAb 2 hrs prior to pneumococcal infection. Following this, mice were intranasally infected with 5x10^6^ CFUs of pneumococcal strain WU2 as described above. On days 3, 7, and 11 post infection, mice were intraperitoneally injected with 25 µg of RB6-8C5 antibody. Mice were monitored as above for the prophylaxis studies.

Macrophages were depleted in mice as previously described (28). 5-7 week old C57BL/6 mice were intraperitoneally injected with 2 mg of clodronate liposome (Encapsula NanoSciences) or control liposomes 2 days prior to infection. Mice were treated with mAb and infected as above, and on days 1, 3, and 6 post infection mice were intraperitoneally injected with 0.5 mg of clodronate liposome or control liposomes. Mice were monitored as above for the prophylaxis studies.

To verify neutrophil and macrophage depletion, a second set of mice treated with RB6-8C5 (14- 5931-86 Invitrogen) (or control mAb) or clodronate liposomes (or empty liposomes) (CLD-8901 Encapsula Nano Sciences) were utilized. Spleens were collected from these mice and cut into small pieces and homogenized into a single-cell suspension through a 40 µm nylon mesh (Corning). The single-cell suspension was centrifuged at 300xg for 10 min at 4 °C, resuspended in 5 mL of 1x RBC lysis buffer for 5 min, and washed twice in sterile PBS buffer. Cells were Fc blocked (#564220 BD Biosciences) for 30 min, the cells were then stained with antibodies specific for CD11b (#60-0112-U025 Tonbo biosciences) and Ly6G (#127625 Biolegend) (for neutrophils) or F4/80 (#157304 Biolegend) and CD11b (for macrophages) for 30 min at 4 °C. The stained cells were washed by resuspending in PBS buffer and centrifuged at 300xg for 10 min at 4 °C. The stained cells were then resuspended in 1 mL of PBS buffer. Myeloid leukocytes were gated among the CD11b+ cells after excluding dead cells positive for staining with live/dead Zombie Aqua dye (#423102 Biolegend). Neutrophils were identified as CD11b+/Ly6G+. Macrophages were recognized as F4/80+ in the CD11b+ population.

Complement depletion in mice was completed as previously described (28). 5-7 week old C57BL/6 mice (Charles River) were intraperitoneally injected with 0.01 U/g of cobra venom factor one day prior to pneumococcal infection, and mice were intraperitoneally injected with 15 mg/kg of mAb 2 hrs prior to pneumococcal infection. Mice were infected with strain WU2 as above, and on days 1,3, 6, 9, and 12 post infection mice were intraperitoneally injected with 0.01 U/g of cobra venom factor. Mice were monitored as above for the prophylaxis studies. To verify our complement depletion procedure, a second set of mice treated with CVF or PBS were utilized. Serum was collected from depleted and non-depleted mice at several time points. Murine serum C3 levels were determined as previously described (29). In brief serum from complement depleted and non-depleted mice was used to coat 96-well plates overnight. The plates were then washed 1x with PBS containing 0.05% Tween-20 (PBS-T) and blocked for 1 hr at room temperature with blocking buffer (2% nonfat milk, 2% goat serum in 0.05% PBS-T). Plates were washed 3x with PBS-T. Following washing a 1:2000 dilution goat anti-mouse C3 antibody-HRP conjugated (#GC3-90P-Z, ICL, Inc) in blocking buffer was added and incubated for 1 hr at room temperature. Plates were washed 3x PBST followed by the addition of TMB, and the reaction was stopped with 2 M sulfuric acid absorbance was measured at 450nm.

### Bacterial burden studies

For primary pneumococcal infection studies, 5-7 week old C57BL/6 mice (Charles River) were intranasally infected with pneumococcal serotype 3 strain WU2 by anesthetizing mice via inhalation of 5% isoflurane and administering 40 µL of 5x10^6^ colony-forming units (CFUs) of bacteria in phosphate buffered saline (Corning). Two hours prior to infection, mice were intraperitoneally inoculated with 15 mg/kg of antibody treatment. On day 3 post infection mice were euthanized, and blood and lungs were collected. Blood was serially diluted and plated to determine bacterial titers. Lungs were homogenized in 1 mL of PBS, and homogenates were then serially diluted and plated to determine bacterial titers as above.

### PhtE cross reactivity binding ELISA

To determine anti-PhtD mAb cross-reactivity with PhtE, we cloned the gene for PhtE from the genome of S. pneumoniae strain TCH8431 (serotype 19A) using primers 5’- ggagatataccatggctAAATTTAGTAAAAAATATATAGCAGCTG-3’ and 3’- gtggtggtgctcgagCGCTATGAGATCAGATAAATTC-5’ via polymerase incomplete primer extension cloning, and the gene was cloned into pET28a. The sequence of the constructed plasmid was confirmed by sequencing and then transformed into *E. coli* BL21(DE3) for protein expression. Single colonies of transformed *E. coli* were picked and cultured in 5 mL of LB medium supplemented with 50 μg/ml kanamycin overnight in a shaking incubator at 37 °C. The overnight culture was then expanded at a 1:100 ratio in 2× yeast tryptone (YT) medium with antibiotic and cultured at 37 °C. After the optical density at 600 nm (OD600) reached 0.5 to 0.7, the culture was induced with 50 μM isopropyl-d-thiogalactopyranoside for 12 to 16 hrs at room temperature. Bacteria pellets were collected by centrifugation at 6,000 × g for 10 min and frozen at −80 °C. Thawed E. coli pellets were resuspended in 10 ml of buffer containing 20 mM Tris (pH 7.5) and 500 mM NaCl and then lysed by sonication. Cell lysates were centrifuged at 12,000 × g for 30 min, and the supernatant was subsequently used for protein purification through a HisTrap column (His-tagged full-length proteins; GE Healthcare) following the manufacturer’s protocols. For recombinant-protein capture ELISAs, 384-well plates were treated with 2 μg/ml of antigen in PBS for 1 h at 37 °C or overnight at 4 °C. Following this, plates were washed once with distilled water before blocking for 1 hr with 2% nonfat milk–2% goat serum in 0.05% PBS-Tween (PBS-T) (blocking buffer). Plates were washed with water three times before serially diluted primary MAbs in PBS were applied for 1 hr. Following this, plates were washed with water three times before application of 25 μL of secondary antibody (goat anti-human IgG Fc; Meridian Life Science) at a dilution of 1:4,000 in blocking solution. After incubation for 1 h, the plates were washed five times with PBS-T, and 25 μL of a PNPP (p-nitrophenyl phosphate) solution (1 mg/ml PNPP in 1 M Tris base) was added to each well. The plates were incubated at room temperature for 1 h before reading the optical density at 405 nm on a BioTek plate reader. Binding assay data were analyzed in GraphPad Prism using a nonlinear regression curve fit and the log(agonist)-versus-response function.

### Flow binding experiments

The ability of mAbs to bind antigen exposed on the surface of *S. pneumoniae* in the presence of the CPS was determined by flow cytometry. Bacteria (1x10^7^ CFUs) were incubated with 10 µg/mL of mAb targeting the capsule and mAb PhtD3 for 30 min at 37 °C. Bacteria were washed with Hanks’ balanced salt solution (HBSS) containing 1% BSA to remove excess stain. Following this, bacteria were incubated with 10 µg/mL of AlexaFlour 488 IgG2a anti-mouse and AlexaFlour 594 anti-human IgG (Invitrogen) antibodies for 30 min at 37 °C. Bacteria were washed twice with HBSS plus 1% BSA and live bacteria were analyzed on a NovoCyte Quanteon flow cytometer. mAb 19F80 was generated using PBMCs obtained from a human subject who received Prevnar-13. PBMCs obtained 28 days after vaccination were stimulated on a NIH 3T3 feeder layer expressing human CD40L, human IL-21, and human BAFF, and screened by ELISA after six days with pneumococcal capsular polysaccharide 19F (ATCC 99-X). The mAb was then generated using human hybridoma technology following single cell sorting as previously described (30). The serotype 4-specific antibody was obtained from Statens Serum Institute and the antibody was labeled with Alexa-488 (anti-S4-A488) following the manufacturer recommendations (Molecular Probes) (31).

### IFA experiments

Encapsulated *S. pneumoniae* TIGR4 bacteria were incubated with 25 µg/mL of anti-S4-A488 or 100 µg/mL of mAb PhtD3 for 60 min at room temperature. TIGR4 bacteria were then washed three times with PBS. Pneumococci labeled with anti-S4-A488 were dropped on a microscope slide and air dried while TIGR4 bacteria incubated with the mAb PhtD3 were incubated with a species-specific secondary antibody labeled with Alexa-488 (Southern scientific) for 60 min at room temperature. Following two washes with PBS these pneumococci were spotted on a microscope slide and air dried. Preparations were mounted with ProLong diamond antifade mountant with DAPI (Molecular Probes). Confocal images were obtained using a Nikon AX R confocal microscope and analyzed with ImageJ version 2.1.051 (National Institutes of Health, USA).

### Co-infection studies

To establish the model of pneumococcal co-infection with influenza virus, 5–7-week-old C57BL/6 mice (Charles River) were anesthetized by inhalation of 5% isoflurane and intranasally challenged with 40 µL of 10^5^, 10^4^, 10^3^, 10^2^ or 0.5x10^2^ focus forming units (FFU) of H1N1 A/California/07/2009 in PBS. For coinfection studies, 5-7 weeks old C57BL/6 mice (Charles River) were anesthetized by inhalation of 5% isoflurane and intranasally challenged with 40 µL of 100 FFU of H1N1 A/California/07/2009 in PBS. After 7 days, mice were anesthetized by inhalation of 5% isoflurane and intranasally challenged with 40 µL of 1x10^4^ CFUs of pneumococcal strain WU2 in PBS. Two hours prior to bacterial challenge, mice were intraperitoneally inoculated with 15 mg/kg of mAb. Mice were weighed and assessed daily and were euthanized when >30% of pre-infection body weight was lost, were nonresponsive to manual stimulation, and/or were exhibiting respiratory distress.

## Results

### Determining the mechanism of protection for mAb PhtD3

While we have previously demonstrated protection by anti-PhtD human mAbs in the mouse model (26), their mechanism of protection *in vivo* remains unknown. Other studies have shown that PhtD mouse mAbs function through macrophage and complement dependent mechanisms (28). To determine if this mechanism is similar for human mAbs we conducted a series of immune depletion experiments and using mAb PhtD3. A key mechanism in defense against pneumococcal infection is phagocytosis, with neutrophils and macrophages being the main mediators. To determine the role of each of these cell types in PhtD3 mediated protection, we tested the efficacy of mAb PhtD3 in immune cell depletion models. First, we depleted neutrophils using an anti-Ly6G mAb delivered two days prior to mAb PhtD3 treatment and pneumococcal infection, and again on days 3, 7, and 11 post infection. No significant difference in survival was seen between mAb PhtD3 vs mAb PhtD3+anti-Ly6G treated groups (90% vs 90%), while significant protection in mAb PhtD3 treated groups was observed compared to isotype control mAb treated groups as previously demonstrated (26) **(Figure 1A)**. To confirm neutrophil depletion in this model, we assessed Ly6G+/CD11b+ cells in spleenocytes of an additional six mice, three of which were given anti-Ly6G mAb or isotype control mAb. The percentage of Ly6G+/CD11b+ cells was reduced in the anti-Ly6G treated mice compared to the isotype control mAb treated mice **(Figure 1B, 1C).**

**Figure 1.**
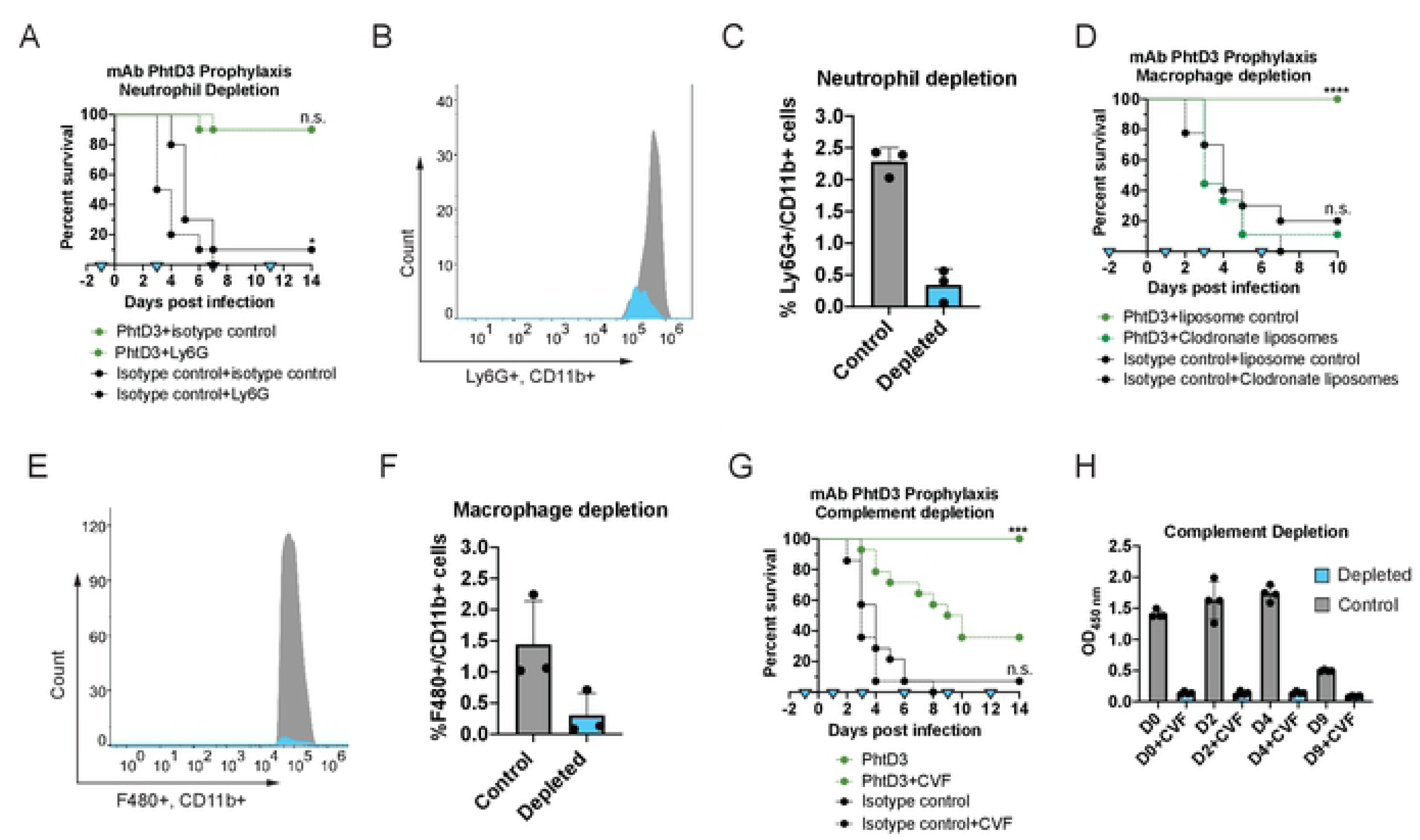
Mechanism of protection by mAb PhtD3. (A) Neutrophil depletion does not diminish protection by mAb PhtD3 in an intranasal infection model of pneumococcal serotype 3 (strain WU2) in 6-8 week old C57BU6 mice. *P=0.0237 via log-rank (Mantel-Cox) test compared to the isotype control+Ly6G group, n.s.=not significant, n=10 mice/group. Blue triangles denotes the days of depletion treatment. (B) Example histogram of Ly6G+/CD11b+ cell population counts. Grey:control animals, Blue: depleted animals. Representative of 1 animal each. (C) Comparison of %Ly6G+/CD11b+ cells Grey: control animals, Blue: depleted animals. n=3 mice/group. (D) Macrophage depletion abolishes protection by mAb PhtD3 in an intranasal infection model of pneumococcal serotype 3 {strain WU2) in 6-8 week old male C57BU6 mice, ****P<0 .0001 compared to the PhtD3+Clodronate liposome treated group; n.s. compared to the isotype control+clodronate liposome treated group. Analysis conducted via log-rank (Mantel-Cox) test. n=9-10 mice/group.Blue triangles denote the days of depletion treatment. (E) Example histogram of F480+/CD1 1b+ cell population counts. Grey: control animals, Blue: depleted animals. Representative of 1animal each. (F) Comparison of o/oF480+/CD 11b+ cells Grey: controlanimals, Blue: depleted animals. n=3 mice/group. (G) Complement depletion partially diminishes protection by mAb PhtD3 in an intranasal infection model of pneumococcal serotype 3 (strain WU2) in 6-8 week old male C57BU6 mice. - P=0.0003 compared to the PhtD+CVF group; ns compared to the isotype control mAb+CVF group.Analysis completed via log-rank (Mantel-Cox) test. n=14 mice/group. Blue triangles denote days of depletion treatment. (H) Serum C3 complement levels in CVFand PBS treated mice. Grey: control animals, Blue:depleted animals. n=3 mice/group

To determine if macrophages are important for mAb PhtD3 efficacy, we tested mAb PhtD3 in a depletion model utilizing clodronate liposomes. Mice were injected with clodronate liposomes to deplete macrophages or empty liposomes as a control two days prior to mAb PhtD3 treatment and pneumococcal infection, and again on days 1, 3, and 6 post infection. Mice treated with mAb PhtD3+empty liposomes demonstrated significantly higher survival when compared to mice treated with mAb PhtD3+clodronate liposome (100% vs 10%) **(Figure 1D)**. Our depletion model was confirmed by flow cytometry using a separate set of mice in which spleenocytes were stained and assessed for the percentage of F480+, CD11b+ cells, which were reduced in the clodronate liposome treated mice **(Figure 1E, 1F)**. This data demonstrated that macrophages play a key role in the protective effects seen with mAb PhtD3 treatment.

To resolve the role of complement in mediating protection by mAb PhtD3, we assessed how depletion of complement affects protection induced by mAb PhtD3 in the mouse infection model. Mice were injected with cobra venom factor (CVF) to deplete complement before administration of mAb PhtD3 and pneumococcal infection, and again on days 1, 3, 6, 9, and 12 days post infection. Mice treated with CVF but not mAb PhtD3 were the first to succumb to infection demonstrating 0% survival. Mice treated with PhtD3+CVF demonstrated prolonged survival compared to isotype control mAb+CVF treated mice. However, treatment with mAb PhtD3+PBS demonstrated significantly increases survival (100% vs 35%) compared to mAb PhtD3+CVF treated mice **(Figure 1G)**. To verify complement depletion, we utilized a separate set of mice treated with CVF or PBS, and utilized an ELISA to measure the levels of C3 present in the serum of these mice at various timepoints, which indicates our treatment regimen successfully depleted C3 **(Figure 1H).** Overall, these data indicate the efficacy of mAb PhtD3 is dependent on macrophages and complement as previously described for anti-PhtD mouse mAbs (28).

To determine what role mAb PhtD3 treatment has on mouse bacterial burden, mice were treated with mAb PhtD3 or isotype control mAb 2 hours prior to infection, and then sacrificed on day 3 post infection for assessment of bacterial titers in the lungs and blood. Mice treated with PhtD3 exhibited lower lung titers than the isotype control group (2.19 vs 6.39 log10 CFU/mL) **(Figure 2A)**. This reduction was also seen in the blood where no bacteria was recovered in the blood of mAb PhtD3 treated mice while bacteria was recovered in the isotype control group (0 vs 2.39 log10 CFU/mL) **(Figure 2A)**. These data suggest the mAb PhtD3 limits the spread of bacteria from the lungs to the blood as a potential mechanism of protection. The protein PhtD is a member of the pneumococcal histidine triad family which has four described members PhtA, PhtB, PhtD, PhtE (32). Since PhtD and PhtE are known to be protective antigens in the mouse model, we tested if our previously discovered PhtD mAbs (26) had cross reactivity with PhtE, which shares 35% sequence identity with PhtD (32). mAb PhtD3 was able to bind to PhtE along with mAb PhtD6, while mAbs PhtD7 and PhtD8 did not bind to PhtE **(Figure 2B)**. Our previous work demonstrated that mAbs PhtD3 and PhtD6 bind to a similar region on the N-terminal region of PhtD. This N-terminal region of PhtD is more conserved with PhtE compared to the C-terminal region.

**Figure 2.**
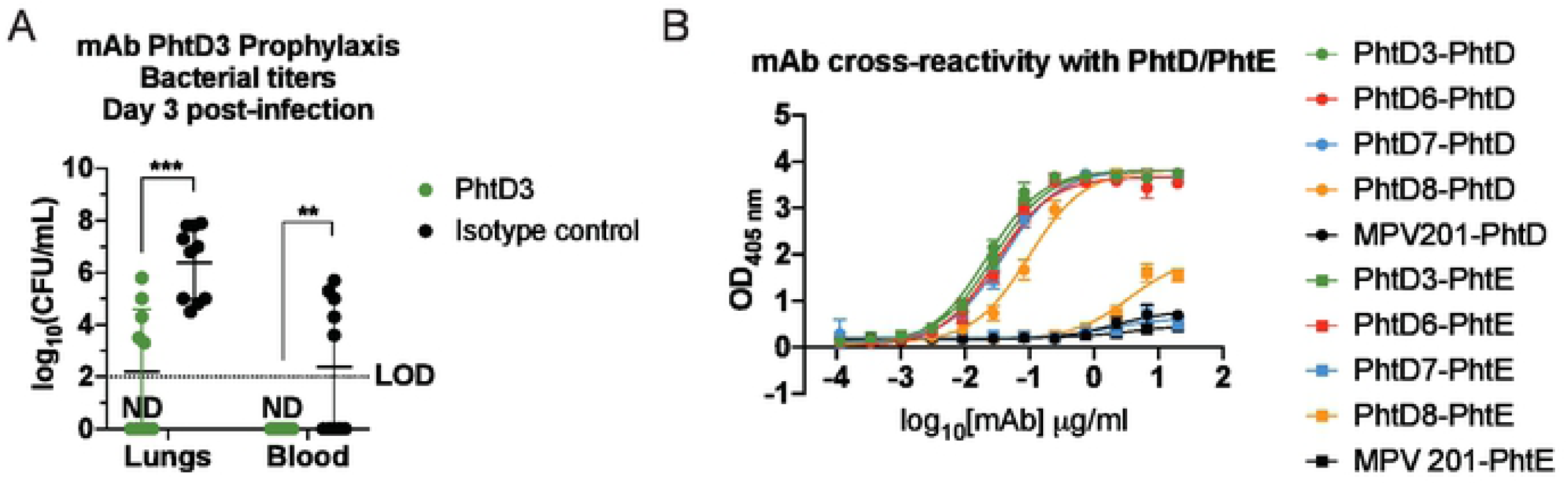
Bacterial burden reduction and cross-reactivity of PhtD mAbs. (A) mAb PhtD3 reduces lung and blood bacterial titers, and binds to family member PhtE.Bacterial burden assessed three days after intranasal pneumococcal infection and prophylactic PhtD3 administration, n=10 mice/group,***P=0.0002;**P=0.009;analysis completed by two-way ANOVA. (B) ELISA binding curves of PhtD mAbs against recombinant PhtD and PhtE proteins.Error bars are the standard deviation of four replicated.

### Assessment of mAb PhtD3 binding in the presence of capsular polysaccharide

One of the most important virulence factors of *S. pneumoniae* is the CPS, which provides protection for the bacteria against the host immune system. However, it is unclear if antibodies are able to access surface proteins in the presence of the CPS. Utilizing a flow cytometric approach, we incubated bacteria with anti-capsule antibodies and mAb PhtD3. We utilized capsule-specific mAbs (19F80 for serotype 19A, and anti-S4-A488 (31) for serotype 4) and co-incubated these antibodies with mAb PhtD3 for pneumococcal strains TCH8431 (serotype 19A) or TIGR4 (serotype 4). We observed binding of both mAb PhtD3 and the capsule-specific antibodies via flow cytometry by gating on double positive events **(Figure 3A)**. The level of double positive events was >90% for serotype 19A, and approximately 30% for TIGR4, suggesting either different levels of PhtD accessibility among pneumococcal serotypes or differing levels of mAb PhtD3 recognition of serotype 19A and 4 PhtD proteins. These data indicate that mAb PhtD3 can bind to encapsulated bacteria for both serotypes 19A and 4. We attempted this study using a previously described anti-serotype 3 mAb (33) and pneumococcal serotype 3 strain WU2, however, we could not obtain consistent binding of this mAb to bacteria likely due to capsule shedding upon mAb binding as previously described (34), since the capsule of pneumococcal serotype 3 is not covalently attached to the surface. However, we have previously demonstrated binding of mAb PhtD3 to serotype 3 bacteria, although it is unclear if the capsule is present or not (26). To verify our findings, we conducted another experiment with pneumococcal serotype 4 (TIGR4), in which bacteria were incubated with mAb PhtD3 followed by a secondary stain labeled with Alexa-488 or with an anti-serotype 4 capsule labeled with Alexa 488 (anti-S4-A488) in addition to staining DNA with DAPI. Three-dimensional reconstruction of z-stacks obtained with a confocal microscope revealed localized foci of PhtD3 on the surface of encapsulated pneumococci **(Figure 3B, top panels)**. Capsule staining with an anti-S4-A488 confirmed the presence of the capsule on TIGR4 bacteria **(Figure 3B, bottom panels).**

**Figure 3.**
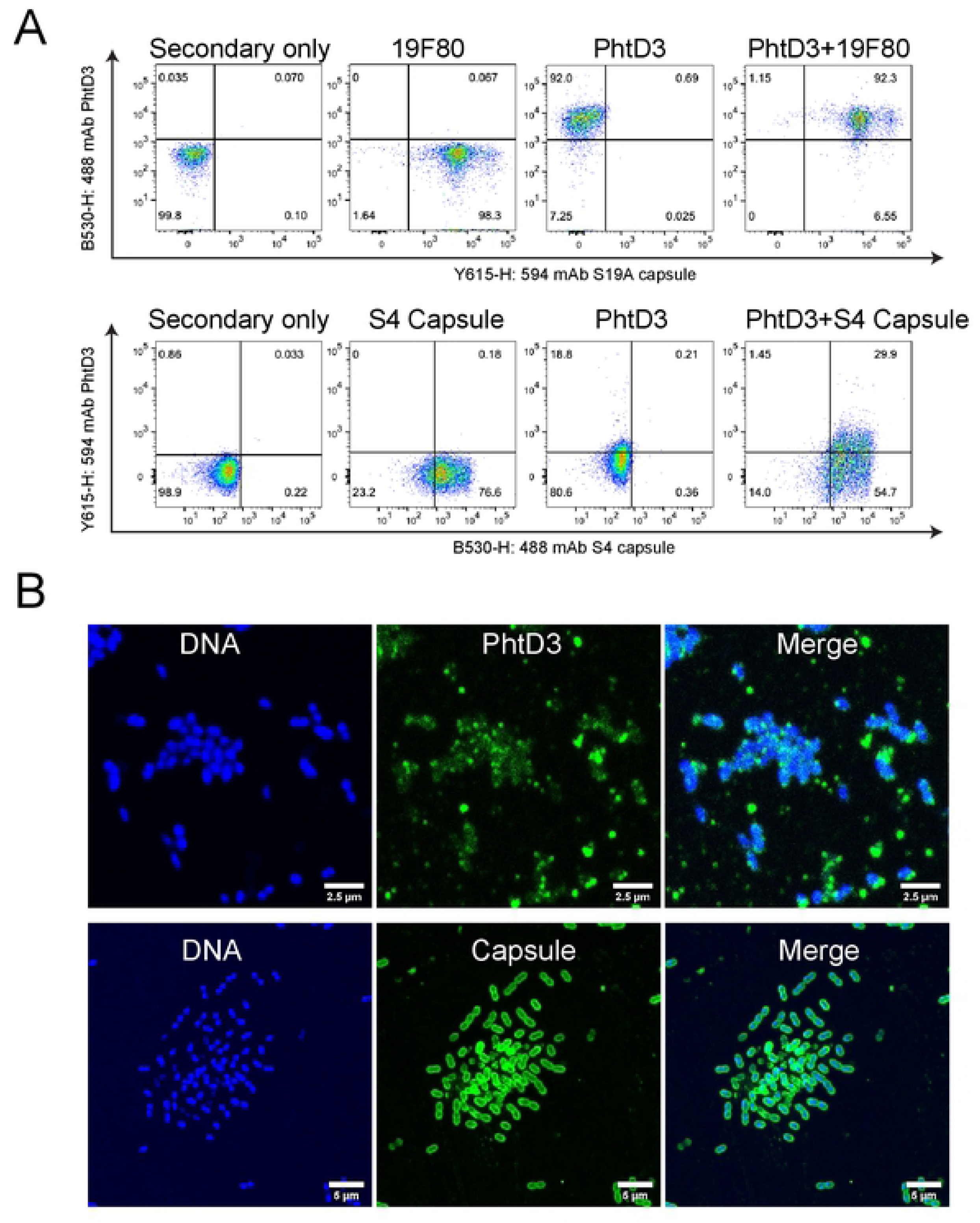
mAb PhtD3 binding to encapsulated bacteria. (A) Flow cytometry analysis of mAb PhtD3 binding to encapsulated bacteria. Serotype 19A bacteria (strain TCH8431) were labeled with mAb PhtD3-lgG2a (with anti-mouse Fe secondary) and/or the human capsule-specific antibody 19F80 (with anti-human Fe secondary). Serotype 4 bacteria (strain TIGR4) were labeled with mAb PhtD3-lgG1 (with anti-human Fe secondary) and/or the rabbit capsule-specific antibody. Figures are representative of one experiment from triplicate repeats. (B) Binding of the anti-PhtD3 mAb to encapsulated pneumococcal serotype 4 (strain TIGR4). Bacteria were stained with an anti­ PhtD3 mAb, followed by a species-specific secondary antibody labeled with Alexa-488 (top panels) or with an anti-serotype 4 capsule labeled with Alexa 488 (bottom panels). The DNA was stained with DAPI. Micrographs were collected with a confocal microscope and projections of -20 xy optical sections (0.1 µm each) are shown.

### Protection against secondary pneumococcal infection

The most clinically useful scenario for the use of anti-pneumococcal mAbs is to treat after symptom onset or once influenza infection is confirmed to protection against secondary pneumococcal infection. Our previous studies demonstrated treatment with mAb PhtD3 24 hours after infection increased survival compared to isotype control mAb treated mice (26). Co-infections with influenza virus and *S. pneumoniae* increase the risk of severe disease and lead to increased rates of hospitalization and mortality (13). To test if mAbs targeting PhtD can protect against secondary pneumococcal infection following influenza infection, we first determined a sublethal dose of H1N1 A/California/07/2009 influenza virus that caused no mortality alone but led to high mortality when combined with a sublethal dose of *S. pneumoniae* using a model previously described (35). Mice were infected with infectious doses of influenza virus of 0.5x10^2^, 10^2^, 10^3^, 10^4^, 10^5^ FFUs. The 10^5^ dose led to 0% survival while the 10^4^ and 10^3^ led to 20% and 60% survival, respectively **(Figure 4A)**. The 10^2^ dose caused no mortality, thus we utilized this dose, similar to previous studies (35), for establishment of the co-infection model. We first infected mice with 10^2^ FFUs of influenza virus, and 7 days later we intranasally infected mice with 4 different doses of *S. pneumoniae* 10^5^, 5x10^4^, 10^4^, 10^3^. The lowest dose of bacteria that led to 100% mortality was 10^4^ **(Figure 4B)**. We then tested the efficacy of mAb PhtD3 in the co-infection model by treating mice 2 hrs prior to pneumococcal infection. The PBS and isotype control mAb treated groups had 0% survival, while mice treated with mAb PhtD3 had delayed mortality and overall survival of 20% **(Figure 4C)**.

**Figure 4.**
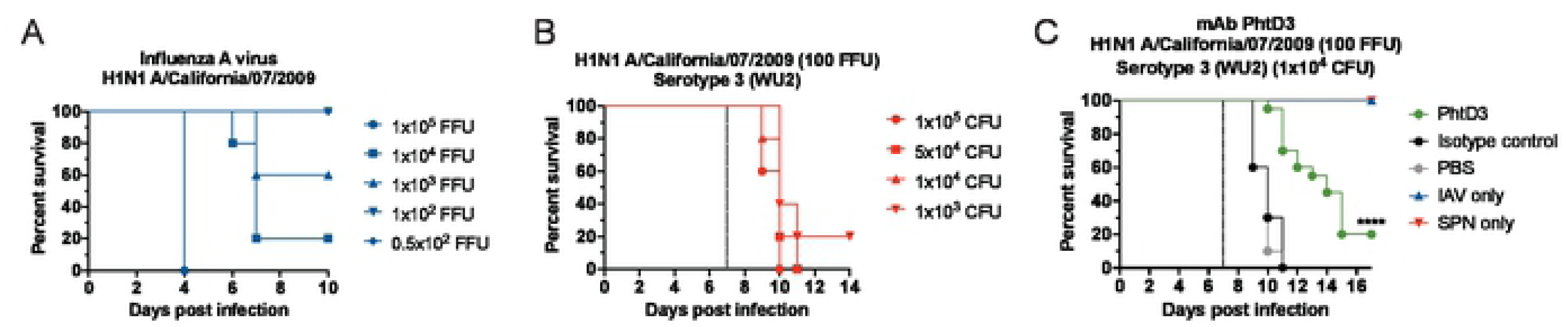
Co-infection with sublethal influenza A virus followed by Spn causes mortality in mice and is improved by PhtD3 mAb treatment. (A) Survival of influenza infection model of influenza A virus. n=5mice/group. (B) Survival of co-infection model of influenza A virus at day O and pneumococcal serotype 3 (strain WU2) at day 7 (dotted line) in 6-8 week old male C57BU6 mice. n=5 mice/group. (C) Protective efficacy of mAb PhtD3 ina co-infection model. Mice were infected with influenza A virus at day O and with WU2 bacteria at day 7 (dotted line). ****P<0.0001 via log- rank (Mantel-Cox) test compared to the isotype control mAb group.Co-infected groups had n = 20 mice/group, while IAV and Spn infected only had n=5 mice/groups.

As the protective efficacy of mAb PhtD3 was much lower in the coinfection model compared to treatment in a model of primary pneumococcal infection **(Figure 1A, 4C)**, we tested the protective efficacy of two additional human mAbs we previously isolated (26) first in a prophylactic treatment model of primary pneumococcal infection as a prelude to the co-infection model. mAb PspA16 targets the N-terminal region of PspA, and while this mAb was shown to have limited serotype breadth, binding to pneumococcal serotype 3 was previously observed (26). Additionally, we included mAb PhtD7, which also targets PhtD and is broadly-reactive, yet binds a conformational epitope located on the C-terminal region of PhtD that does not overlap with mAb PhtD3 (26). PspA16 was able to significantly prolong survival of mice compared to isotype control mAb and PBS treated mice, 50% vs 0% vs 0% respectively **(Figure 5A)**. Mice treated with mAb PhtD7 demonstrated significantly higher survival 90% vs 10% compared to isotype control mAb treated mice **(Figure 5B)**. As mAb PhtD7 demonstrated robust efficacy in a prophylactic model compared to mAb PspA16, we tested mAb PhtD7 in a treatment model where we administered the mAb 24 hours post infection as previously described for mAb PhtD3(26). mAb PhtD7 treated mice were rescued from pneumococcal infection compared to isotype control mAb treated mice (100% vs 20%) **(Figure 5C)**. We next tested the efficacy of mAb PhtD7 in the co-infection model, yet we observed limited survival for mAb PhtD7 treated mice compared to isotype control mAb treated mice (15% vs 0%), similar to that observed for mAb PhtD3 **(Figure 5D).**

**Figure 5.**
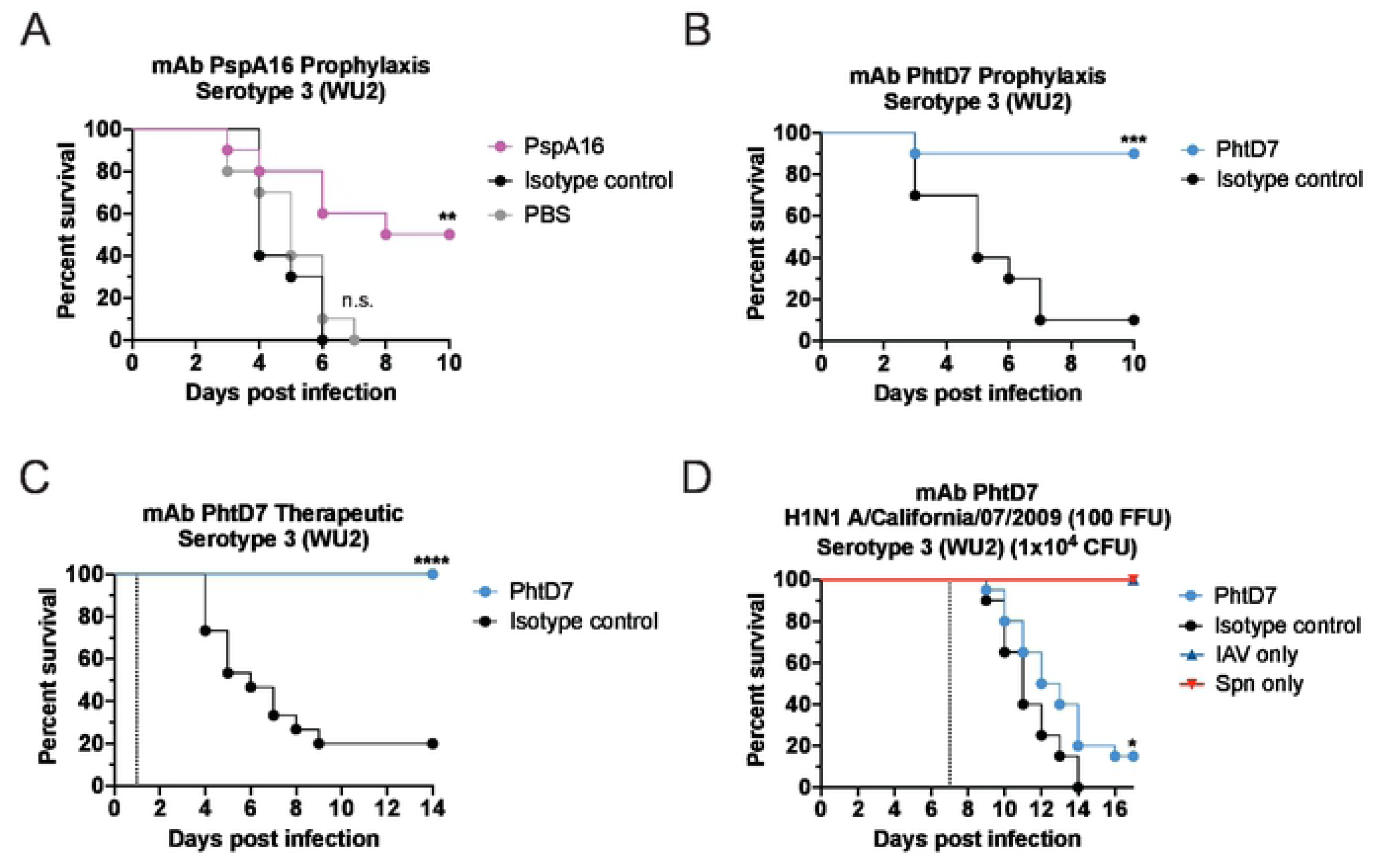
Protective efficacy of mAbs PspA16 and PhtD7. (A) Prophylactic efficacy of mAb PspA 16 in an intranasal infection model of pneumococcal serotype 3 (strain WU2) in 6-8 week old male C57BU6 mice. **P=0.001; ns, not significant via log-rank (Mantel-Cox) test compared to the isotype controlgroup. n=10 mice/group. (B) Prophylactic efficacy of PhtD7 in an intranasal infection model of pneumococcal serotype 3 (strain WU2) in 6-8 week old male C57BL/6 mice. ***P=0.0006; via log-rank (Mantel-Cox) test compared to the isotype control group. n=10 mice/group. (C) Treatment efficacy of mAb PhtD7 in an intranasal infection model of pneumococcal serotype 3 (strain WU2) in 6-8 week old male C57BU6 mice. ****P<0.0001;via log-rank (Mantel-Cox) test compared to the isotype control group. n= 15 mice/group. (0) Protective efficacy of mAb PhtD7 in a co-infection model. Mice were infected with influenza A virus at day O and with WU2 bacteria at day 7 (dotted line). *P=0.0264 via log-rank (Mantel-Cox)

As treatment with mAbs PhtD3 and PhtD7 is highly effective in the mouse model of primary pneumococcal infection, both prophylactically and therapeutically, yet protection was reduced in the co-infection model, we next tested the hypothesis that combining two mAbs that bind distinct nonoverlapping epitopes could lead to a synergistic effect in protecting against secondary pneumococcal infection. In a first experiment, we tested the therapeutic efficacy of combining mAbs PhtD3 and PhtD7 in a model of primary pneumococcal infection. In these experiments, the individual mAb dose was halved (7.5 mg/kg) compared to previous experiments so that the total mAb dose was kept consistent (15 mg/kg). Mice treated with both mAbs PhtD3 and PhtD7 24 hours post pneumococcal infection has significantly improved survival compared to mice treated with an isotype control mAb (85% vs 10%) **(Figure 6A)**. Based on these data, we next utilized this combination in our co-infection model. We discovered that the combination of both mAbs PhtD3 and PhtD7 led to a synergistic effect resulting in a robust increase in survival compared to the isotype control mAb treated mice 85% vs 0% respectively **(Figure 6B)**.

**Figure 6.**
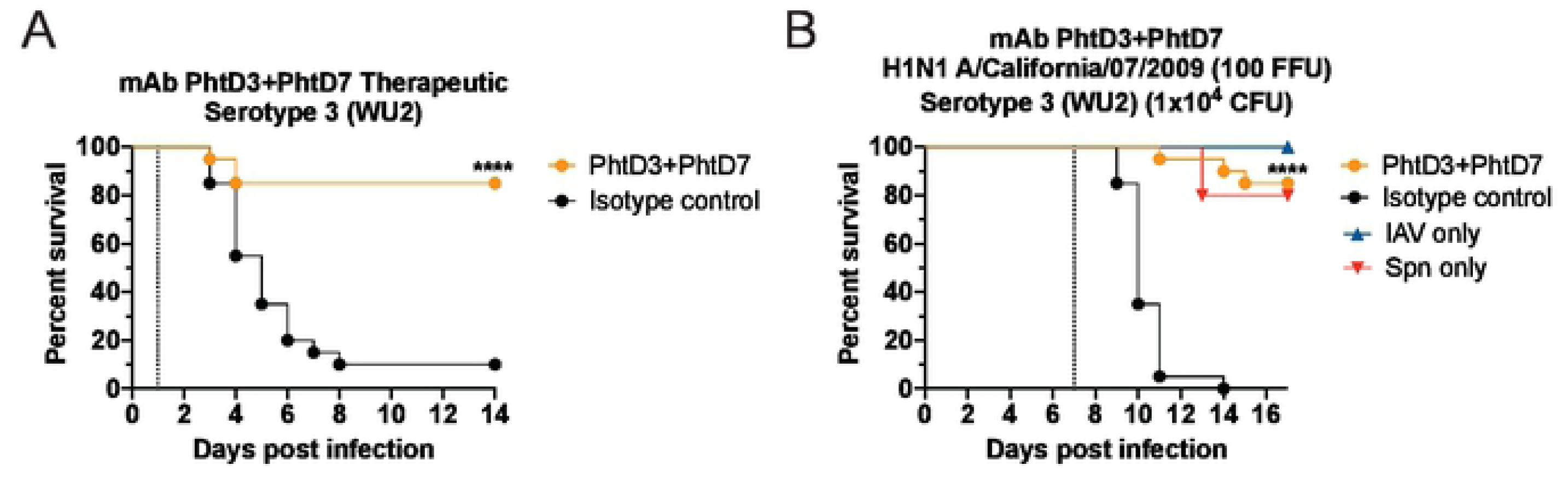
Synergistic effects of mAb PhtD3 and mAb PhtD7 co-administration. (A) Treatment efficacy of mAbs PhtD3+PhtD7 in an intranasal infection model of pneumococcalserotype 3 (strain WU2) in 6-8 week old male C57BU6 mice. ****P<0.0001 via log-rank (Mantel-Cox) test compared to the isotype controlgroup. n = 20 mice/group. (B) Protective efficacy of mAb PhtD3 and PhtD7 in a co-infection model. Mice were infected with influenza A virus at day O and with WU2 bacteria at day 7 (dotted line). ****P<0.0001;via log-rank (Mantel-Cox) test compared to the isotype control group.Co-infected n = 20 mice/group, IAV and Spn infected only n=5 mice/group.

## Discussion

In this study, we demonstrated that the protective efficacy of the anti-pneumococcal human mAb PhtD3 is dependent on macrophages and the complement system, but not neutrophils. Additionally, we demonstrated the protective efficacy of a combination therapy utilizing mAb PhtD3 and mAb PhtD7 in the context of a co-infection model with a primary IAV infection and a secondary pneumococcal infection. The pneumococcal protein PhtD has previously been explored as a vaccine candidate however human trials have failed to elicit protection in pneumococcal carriage in infants (36, 37). A previous study indicated that PhtD mAbs function via complement and macrophage mediated protection (28). These findings were confirmed for the human derived mAb PhtD3, and is likely similar for mAb PhtD7 and additional anti-PhtD mAbs we previously isolated (26).

One of the key phagocytes in protection against pneumococcal infection are neutrophils. Neutrophils phagocytose and degrade *S. pneumoniae* via neutrophil elastase and cathepsin G (38), and previous depletion studies have shown the importance of neutrophils in bacterial clearance (39). Depletion of neutrophils led to no change in the survival of mAb PhtD3 treated mice compared to nondepleted PhtD3 treated mice. In addition, macrophages play a major role in protection against pneumococcal infection, and upon depletion of macrophages the efficacy of mAb PhtD3 was lost. The essential role of macrophages has been demonstrated previously in the protective efficacy of other mAbs targeting PcpA (28) and the CPS (40). Mice depleted of complement prior to infection had reduced survival when treated with mAb PhtD3, suggesting that although not completely necessary for mice survival, complement likely enhances the ability of macrophages to phagocytose mAb PhtD3-coated bacteria. To further elucidate the mechanism of protection for mAb PhtD3, we measured lung and blood bacterial titers at day 3 post infection and observed a significant reduction in lung bacterial titers with half the mAb PhtD3-treated mice demonstrating bacterial titers below our limit of detection. Additionally, we were unable to recover any bacteria from the blood of PhtD3 mice while we were able to collect bacteria for the isotype control mAb mice. These data suggest mAb PhtD3 treatment limits spread of bacteria in the lungs to the blood. The ability of mAb PhtD3 to also bind to PhtE is quite fascinating as PhtD and PhtE only share 35% sequence identity. This phenomenon of binding to two antigens of the same family likely allows for more mAbs to bind to the bacteria due to more available targets.

A key component of pneumococcal virulence is the capsule polysaccharide. Although we have previously described broad binding of four anti-PhtD human mAbs to several pneumococcal serotypes (26), we did not confirm this binding took place in the presence of the bacterial capsule. We utilized antibodies that target the capsule of the bacteria in addition to mAb PhtD3 to demonstrate that mAb PhtD3 is able to bind to bacteria in the presence of the capsule for serotypes 19A and 4. To our knowledge this is the first example confirming accessibility of protein antigens on encapsulated bacteria.

Bacterial co-infections with viruses represent a considerable challenge to public health, and for *pneumoniae*, influenza virus is the major pathogen leading to secondary pneumococcal infections (13). Antibiotic-resistance further compounds the difficulty of treating co-infected patients (41), so the search for alternative therapies is of significant importance. Influenza virus is able to suppress alveolar macrophage cell function within 7 days of infection (22, 42). Influenza lung damage is usually greatest on day 6 post infection (18), and the greatest susceptibility to secondary bacterial coinfection occurs near day 7 in humans and mice (43). Due to this observation, we utilized a co-infection model that started the secondary bacterial infection 7 days post influenza infection. However, in this model the efficacy of mAb PhtD3 was reduced compared to the protective efficacy observed in primary infection studies. Due to the reduced efficacy of mAb PhtD3, we determined if other mAbs were capable of greater protection. Our first attempt utilized a mAb we previously isolated, mAb PspA16 (26), which targets PspA, another candidate antigen for protein-based vaccines (44–46). Utilizing our prophylactic model, we found that administration of mAb PspA16 led to 50% survival vs 0% in our control groups. Due to the lower survival seen compared to PhtD3 we tested the efficacy of mAb PhtD7, which appears to bind to a unique conformational epitope that is dependent on amino acids 341-838 of PhtD, yet does not bind 341-647 or 645-838 fragments (26). mAb PhtD7 conferred an 80% increase in survival compared to isotype control treated mice. Additionally, mAb PhtD7 had significant protection in a pneumococcal treatment model. We then tested the efficacy of mAb PhtD7 in the influenza/pneumococcal co-infection model, however, mAb PhtD7 displayed reduced efficacy in this model similar to mAb PhtD3.

Previous research with viruses such as Ebola virus, Hepatitis C virus, and SARS-CoV-2 have demonstrated that synergistic protective effects are observed when using a combination of mAbs targeting unique epitopes (47–49). We wanted to determine if we could produce a synergistic protective effect with mAbs binding to two different epitopes for pneumococcal infection. We first administered both mAbs PhtD3 and PhtD7 24 hrs post infection, to determine if we could confirm a protective effect similar to our previous treatment data using only PhtD3 (26), which led to a 75% increase in survival compared to the isotype control mAb group. In the co-infection model, administration of the mAb cocktail led to a robust effect leading to 85% increase in survival compared to the control group.

Previous studies have demonstrated that influenza primary infection ablates alveolar macrophages (22). However, it is clear from this study and others (28) that anti-PhtD mAbs function through a macrophage dependent mechanism of protection. Thus, further exploration of the protective mechanism in the context of co-infections is needed, including additional assessment of PhtD protein epitopes required for protection, and reduction of mAb doses to determine if synergistic mAbs can be used for dose sparing. In summary our research further strengthens the results of our previous studies that human mAbs to conserved surface antigens are viable for prevention and treatment of pneumococcal infections. We were also able to elucidate *in vivo* functions of our mAbs, finding they function through macrophage and complement dependent mechanisms. Additionally, we demonstrate for the first time that anti- pneumococcal human mAbs can protect against secondary pneumococcal infection following influenza virus. Further studies will determine if these mAbs are effective against multi-drug resistant strains of *S. pneumoniae* or against secondary infection with additional respiratory viruses.

## Acknowledgements

We thank the International Reagent Resource for the H1N1 A/California/07/2009 influenza virus. Authors also thank Ana Vidal from the Department of Microbiology and Immunology, University of Mississippi Medical Center, for their assistance in some laboratory assays. This work was funded by the American Lung Association Innovation Award (J.J.M.) and NIH K01OD026569 (J.J.M.). JEV is in partly supported by NIH 1R21AI151571-01A1. F.R. was supported by National Institutes of Health NIGMS grant GM109435, Post-Baccalaureate Training in Infectious Diseases Research.

